# Intrathecal [^64^Cu]Cu-albumin PET reveals age-related decline of cerebrospinal fluid (CSF)-lymphatic efflux

**DOI:** 10.1101/2023.01.18.524645

**Authors:** Azmal Sarker, Minseok Suh, Yoori Choi, Ji Yong Park, Yun-Sang Lee, Dong Soo Lee

## Abstract

Age-related cognitive decline is associated with dysfunctional lymphatic efflux of cerebrospinal fluid (CSF) through meningeal lymphatic vessels. Intrathecal [^64^Cu]Cu-albumin positron emission tomography (PET) was applied in mice for the evaluation of lymphatic efflux of CSF and its age-related variation. [^64^Cu]Cu-albumin PET was done at multiple time points after intrathecal injection of [^64^Cu]Cu-albumin with the infusion speed of 700 nl/min in the adult and aged mice of 15–25 months old age. CSF clearance and paravertebral lymph nodes were quantified after injection and at later stationary phase. Representing perturbed state by 6 μl (1/7 of CSF volume with twice the production rate for 9 minutes of intrathecal injection) and at the next day of stationary return of CSF dynamics in mice, CSF clearance half-time from the subarachnoid space was 93.4 ± 19.7 in adult and 123.3 ± 15.6 minutes in aged mice (p = 0.01). The % injected dose at 4, 6 and 24 hours were higher in aged mice than in the adult mice (p < 0.05) and the visualized paravertebral lymph node activity tended to be lower in the aged, which was different from [^64^Cu]Cu-NOTA or [^64^Cu]Cu-ESION PET. [^64^Cu]Cu-albumin PET enabled quantification of CSF-lymphatic efflux over all the levels of brain spinal cords and visualization with quantifiability of lymph node activity. [^64^Cu]Cu-albumin PET revealed an age-related decrease in CSF-lymphatic efflux due to less efflux from the subarachnoid space, especially at stationary phase in the aged mice.

## INTRODUCTION

After the discovery of the meningeal lymphatics in 2015 (1, 2), interstitial fluid (ISF) – cerebrospinal fluid (CSF) – meningeal lymphatic efflux pathway is considered to be the major route for brain and spinal cord waste disposal. The same pathway is also considered to be the route of leukocytes to enter the CSF and consequently brain parenchyma and spinal cord via pia or ependyma lining the brain and spinal cord (3, 4). Mouse and rat were found to use their dual pathway of efflux via the nasal cavity and meningeal lymphatics for waste disposal (5-7) while humans use mainly the dura lymphatics not using nasal cavity (8-11). ISF pass through glia limitans to perivascular space which is communicating with CSF space so that any waste in ISF come out to lymphatics of dura meningeal lymphatics of brain and spinal cord (12). This was named as glymphatic drainage by Nedergaard (13), function and dysfunction of which was investigated in rodents using fluorescent macromolecules (14) and in humans using gadolinium contrast MRI (15).

Glymphatic/lymphatic imaging was done using contrast administration via cisterna magna in rodents using fluorescent microscopy or MRI (12,15-17) or L4/5 intervertebral intrathecal administration in humans using MRI (8-11). Both of the approaches tried to visualize reflux of CSF to perivascular space for evaluation of glymphatic function and CSF/lymphatic efflux for CSF lymphatic waste drainage. In our preliminary study, intra-cisterna magna administration of [^64^Cu]Cu-albumin were too invasive to disclose subtle changes of CSF-lymphatic efflux kinetic (18). Optimal use of tracer quantity and optimal volume was necessary for evaluation of CSF-lymphatic efflux. Unlike the initial reports to reveal glymphatic function using fluorescence microscopy in rodents (12,15,17,19), recent reports especially using intrathecal MRI contrast agents showed minute differences of submeningeal to perivascular space reflux (20-22) to obliterate the use of intrathecal contrast MRI for evaluation of glymphatic function. In contrast, CSF-lymphatic efflux study using intrathecal contrast MRI and cisternography PET (18) was with highest contrast without background and thus superior for disclosing the primary/secondary sentinel lymph nodes down the skull along the vertebral columns in rodents (21-23).

The contribution of CSF-lymphatic efflux from brain is now known to be mandatory for CNS health because the disposal by efflux is essential for maintaining the homeostasis of brain/spinal cord function incremental to the in situ clearance of debris/waste by brain glial cells especially during slow wave sleep or during anesthesia (reviewed in (24)). Dysfunction beyond the physiologic range of variation might be subtle and delicate, for example, with the aging-related changes of CSF-lymphatic efflux (25-31). Thus, we need to minimize the volume of intrathecal contrast not to perturb the physiological stationarity, however, not too small for CSF circulation to maintain to deliver the administered tracers along the CSF flow, which starts from ventricular choroid plexus via aqueduct/spinal canal, cisterns, while rounding and soaking perivascular spaces, via the entire subarachnoid space. Then CSF passes through arachnoid barrier cell layer of brain and spinal cord to reach meningeal extracellular space eventually enter the lymphatic channels of cranial/vertebral dura (32). Several radionuclide/radiopharmaceuticals, which have been used clinically for radioisotope cisternography for decades, would suit for this CSF-lymphatic efflux imaging. We chose [^64^Cu]Cu-albumin and already established [^64^Cu]Cu-albumin PET protocols for mouse CSF-lymphatic efflux imaging (18). From this preliminary study, [^64^Cu]Cu-albumin PET could be used for visualization of the primary/secondary sentinel lymph nodes down the skull along the vertebral column in mice (18).

In this study, we corroborated our protocol with a simple model of aging in mice and looked for the possible variations of CSF-lymphatic efflux with aging using intrathecal [^64^Cu]Cu-albumin PET by visual and quantitative analysis of CSF clearance, mean and temporal progress after injection, lymph nodes’ retention immediately after injection and later when the bodily physiology came back to stationary phase. Mouse CSF, which replenishes 10 or more times by production per day, was perturbed with 1/7^th^ amount of total volume, 350 μl injected within 9 minutes and then let at rest for a day and then imaged again for assessing the stationary CSF-lymphatic efflux and lymph node influx/efflux. We also checked whether there is any evidence that the intrathecal [^64^Cu]Cu-albumin came out differentially from the cranial CSF and/or spinal cord CSF to reach cervical and/or sacral lymph nodes.

## MATERIALS AND METHODS

### Animals

Male C57BL/6 mice of 2-9 months of age were used as the adult whereas mice of 15 to 25 months of age as the aged (33). Mice were housed 2-5 per cage with free access to standard food and potable water. The housing room was maintained at a constant temperature of 22 - 24 °C with a 12/12-hour light and dark cycle. A Y-maze test was conducted to assess the behavior difference between groups. This study shares an animal cohort with previously published literature (18), and the previous study dealt with the establishment of [^64^Cu]Cu-albumin PET to evaluate CSF-lymphatic efflux in the adult mice.

### [^64^Cu]Cu-albumin radiolabeling

The human serum albumin, a molecular weight of 66.5 KDa, was radiolabeled with [^64^Cu]Cu (t1/2 = 12.7 hr, β^+^ = 655 keV 17.8%, β^‒^ = 579 keV 38.4%) by click chemistry-based method (34). Briefly, 1) ^64^Cu in HCl solution was dried with a nitrogen flow of 5-10%, 2) 1 M sodium acetate buffer was added to adjust the pH to 5.3, 3) NOTA-N_3_-1300 nmol/mL, 10 μL was added and the mixture was placed in a heat block at 60-70ºC for 10 minutes, 4) The labeled NOTA-N_3_ of [^64^Cu]Cu NOTA-N_3_ was added to 1 mg/mL of ADIBO-albumin (Human serum albumin, 66.7 kDa) and then was placed in an orbital shaker for 10 minutes, 6) the mixture was transferred to filter fitted tube followed by centrifugation at 15,000 rpm for 5 minutes. Labeling efficiency of [^64^Cu]Cu-albumin was confirmed to be 99% on instant thin layer chromatography (ITLC) with 0.1 M citric acid solution as a developing solvent.

### Intrathecal [^64^Cu]Cu-albumin injection and PET image acquisition

Intrathecal administration followed the established procedure having been used for preclinical or clinical radioisotope cisternography (18). Briefly, an intrathecal injection was performed after palpation of the L4 spine. A 1 cm-long sagittal incision was made and a needle was inserted through the muscles up to an appropriate length and watching for the tail to flick. [^64^Cu]Cu-albumin injection was done using the syringe pump (Harvard Apparatus) at an infusion speed of 700 nl/min with the total injected volume being 6 μL containing an amount of radioactivity that ranged between 0.74 and 1.85 MBq among the batches. The needle was then kept in place until the first imaging session was over. Then, the needle was removed and the wound was closed.

Genisys PET box (Sofie Biosciences) was used for whole-body PET acquisition of static PET in list mode for 6 minutes at each of the image acquisition time points, which were 9-minutes, 2, 4, 6, 12, and 24 hours post-injection for adult mice, and 9 minutes, 2, 4, 6, 17, and 24-hours post-injection for aged mice. The injection and image acquisition were done under isoflurane anesthesia, 3% for induction, and 2.8% for maintenance with inhalation of oxygen 500 ml/min.

### Visual image analysis and quantitation

Using the MIM software, separate 3D regions of interest (ROI) were drawn and the counts were obtained from the entire subarachnoid space. Subarachnoid space counts of 9-minutes image (h0) was used as a surrogate of the injected dose for normalization of the counts for all the other ROIs at any given time point (h_n_) which was expressed as %ID. The %ID of each time point (h_n_/h_0_) was plotted against the corresponding time points to generate the time-%ID curves. Using a one-compartment exponential model of the GraphPad Prism, half-life (t_1/2_) was calculated for each mouse. The coefficient of determination R^2^ > 0.95 considered to be a good fit. Unpaired t-test with Welch’s correction was used to compare the half-lives between the groups.

### Assessment of in-vivo stability of [^64^Cu]Cu-albumin

CSF were collected from an adult mouse, one and four hours after intrathecal injection of [^64^Cu]Cu-albumin and ITLC were performed per using the routine protocol.

## RESULTS

### Behaviour difference between adult and aged mice

Y-maze test revealed a significant decline (Mann Whitney p < 0.05) in the spatial memory among the aged mice compared with the adult mice (**Supplementary Figure 1)**.

### Visual comparison of intrathecal [^64^Cu]Cu-albumin PET images in adult and aged mice

**Figure 1A** shows a schematic of intrathecal [^64^Cu]Cu-albumin distribution along the subarachnoid space and further drainage through the meningeal lymphatics of brain and spinal cord dura. On maximal intensity projection (MIP) animation image of a mouse (**Supplementary Movie 1**), [^64^Cu]Cu-albumin reach via brain meningeal and spinal epidural lymphatics to the cervical lymph nodes, the pelvic lymph nodes and others and further via the heart and probable systemic circulation to the liver.

**Figure 1.**
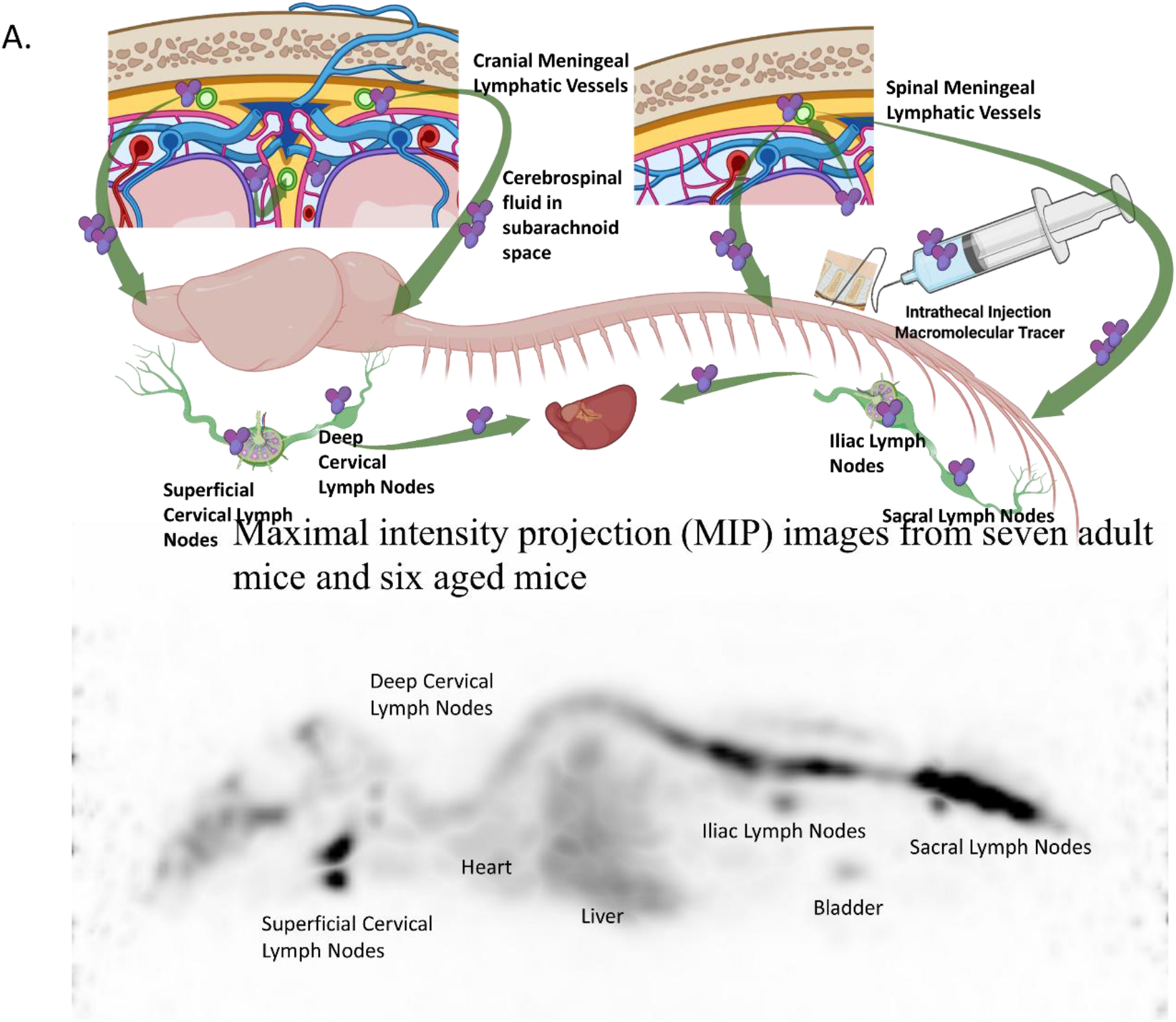
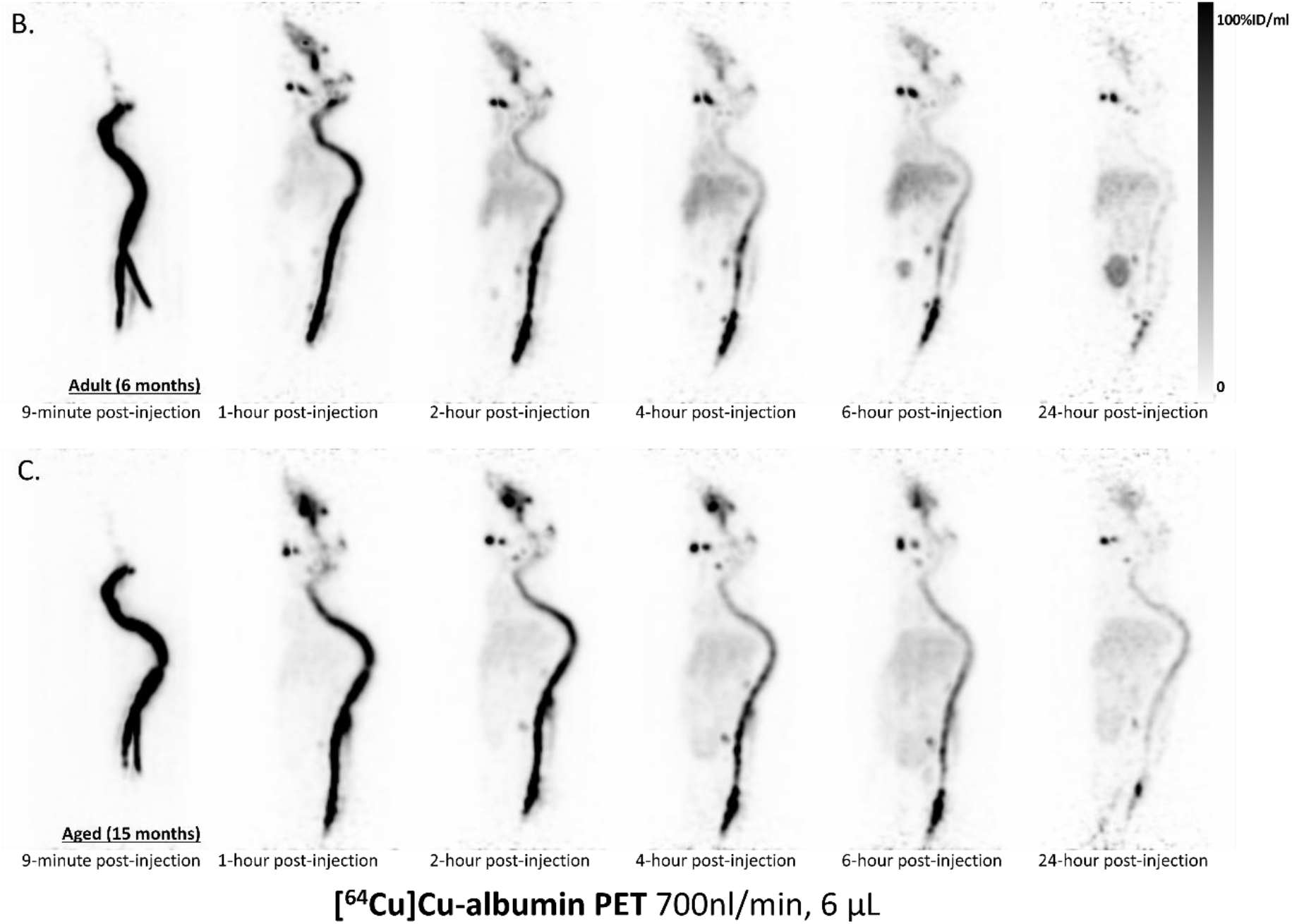
Whole body PET maximum intensity projection images after intrathecal injection of [^64^Cu]Cu-albumin showing the organs involved in the bio-distribution of [^64^Cu]Cu-albumin (A), the temporal change of biodistribution in an adult mouse (B), and an aged mouse (C). The shaded bar on right indicates the activity concentration of the tracer.

MIP animation images from seven adult mice and six aged mice were assessed visually and individual representative images were presented in **Figure 1B-C** and **Supplementary Movie 2**. There was an intense [^64^Cu]Cu-albumin activity in the spinal subarachnoid space with negligible tracer concentration in the cranial and the most caudal parts of the subarachnoid space at the 9-minutes images, which later filled at later images in both the adult and aged mice (**Figure 1B-C**).

On the 1 to 6 hours images in adult mice (**Figure 1B**), radioactivity cleared gradually from the cervical, thoracic and lumbar subarachnoid spaces in adult mice while the hepatic activity continued to increase. The subarachnoid space radioactivity of these adult mice was cleared nearly at 24 hours, with sustained retention in the nasal region, the cervical lymph nodes, the pelvic lymph nodes, and the sacral subarachnoid space. In contrast, on the 1 to 6 hours post-injection in aged mice (**Figure 1C**), radioactivity cleared slowly from the cervical, thoracic and lumbar subarachnoid space than in adult mice. Their hepatic activity continued to increase but intensity was lower than in adult mice.

At 24 hours, the retained activity within the subarachnoid space of the aged mice was higher than that of adult mice. Radioactivity in the nasal region, the cervical lymph nodes, the pelvic lymph nodes, and the sacral subarachnoid space were similar between adult mice and aged mice. Sacral activity was faint in aged mice while the subarachnoid space activity was higher than in adult mice on visual assessment.

### Quantitative comparison of clearance from subarachnoid space of intrathecal [^64^Cu]Cu-albumin between adult and aged mice

The time vs %ID line plot (**Figure 2A**) showed an overall temporal decline in the %ID from the entire subarachnoid space, representing the ‘lymphatic efflux of CSF’ in both the adult and an aged group of mice. The mean clearance halve-lives for adult and aged groups were 93.4 ± 19.7 (range 72.3 – 122.2) and 123.3 ± 15.6 (range 97.9-141.4) minutes, respectively. The mean clearance half-life for aged mice was significantly higher (**Figure 2B**) than that for adult mice (p = 0.01). The %IDs for aged mice at 4, 6, and 24 hours post-injection were also significantly higher (**Figure 2C-G**) than those in adult mice (p < 0.05). Clearance was slower in the aged mice and retention was higher in the subarachnoid space than in the adult mice. The clearance parameters of sub-arachnoid space in individual mice are shown in **Supplementary Table 1**.

**Figure 2.**
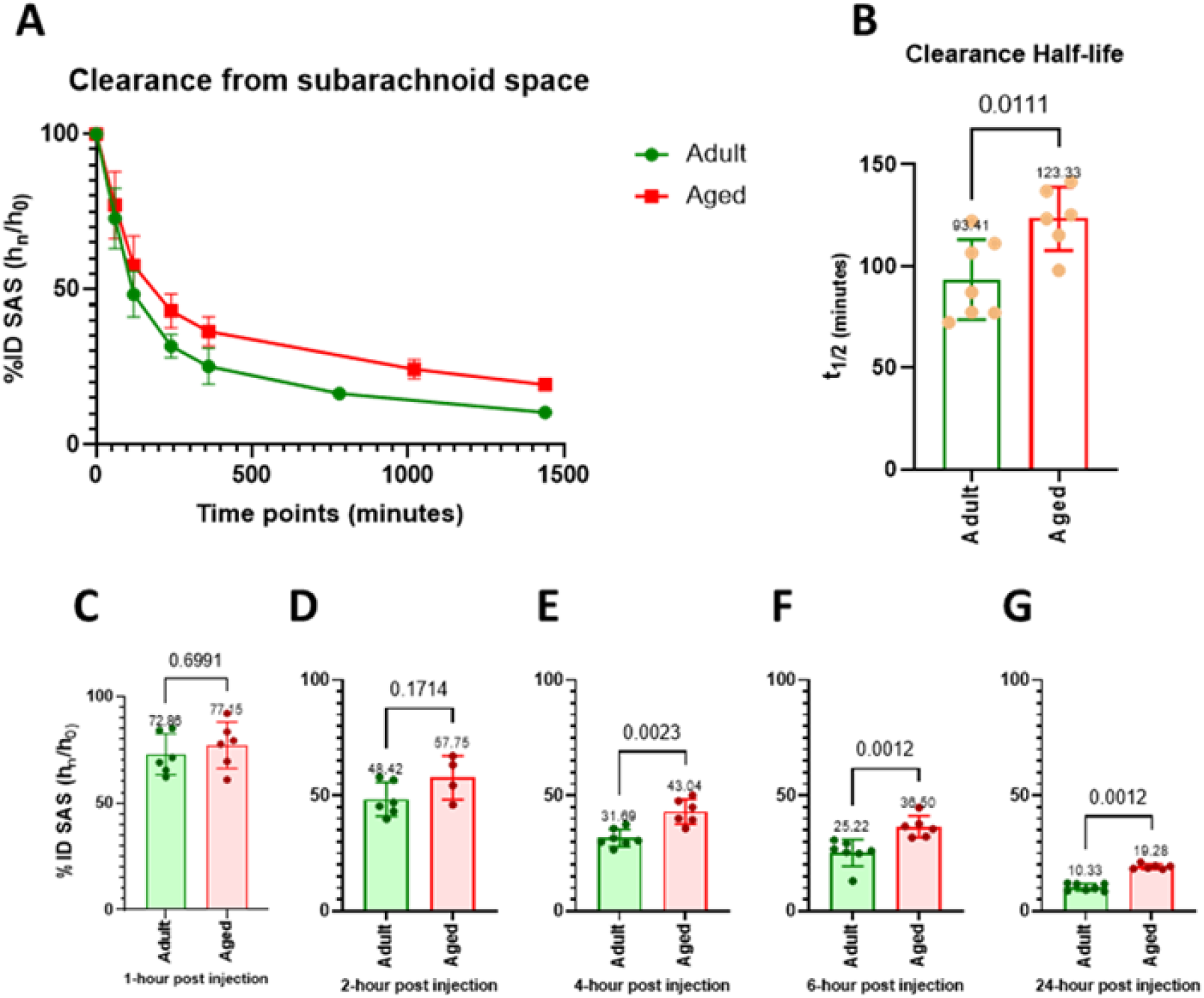
Comparison between adult and aged mice showing time-%ID line-plots for clearance from the entire subarachnoid space (A), bar-plots with pairwise comparison for clearance halve-lives (B), and bar-plots with pairwise comparison for %IDs at the matched time-points (C-G).

### In-vivo stability of subarachnoid space-staying [^64^Cu]Cu-albumin after intrathecal injection

On ITLC, the intact [^64^Cu]Cu-albumin remains peaked at the origin and degraded [^64^Cu]Cu-albumin fragments with variable sizes run to reach the solvent-front to make the second peak representing detached unbound [^64^Cu]Cu-NOTA-N_3_. Single peak was observed at the origin for the CSF sample (**Figure 3A)** to reveal that [^64^Cu]Cu-albumin was in intact form in subarachnoid space at 1 hour and 4 hours after intrathecal injection. Intact [^64^Cu]Cu-albumin of blood and urine samples was peaked at the origin and degraded fragments thereof moved on to the solvent-front at 1 and 4 hours, which amount increase as time lapsed compared with 1 hour **(Figure 3B-C)**.

**Figure 3.**
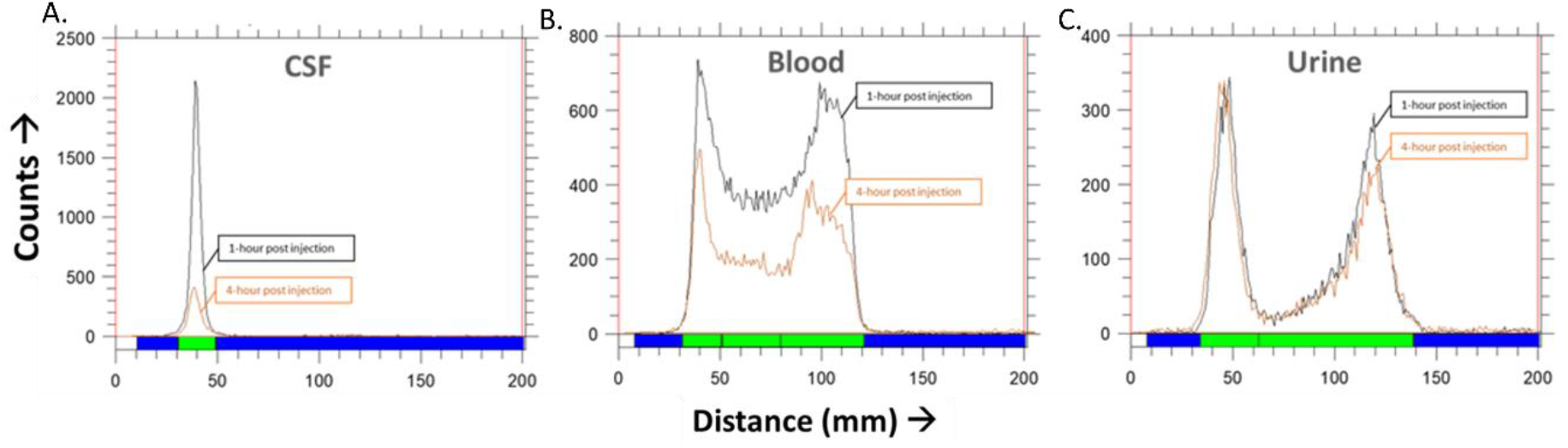
Radioactivity of CSF at 1 hour and 4-hours following the intrathecal injection with chromatograms for the [^64^Cu]Cu-albumin at the corresponding time points show a single peak for the CSF comprising of greater than 90% of the radioactivity at the origin that represented the intact [^64^Cu]Cu-albumin. In contrast, degraded fragments of [^64^Cu]Cu-albumin moved to the solvent front in variable degree for the blood and urine samples 1 or 4 hours after intrathecal [^64^Cu]Cu-albumin injection.

## DISCUSSION

In this study, using [^64^Cu]Cu-albumin PET, we corroborated the CSF-lymphatic efflux of the previous studies by fluorescent or MRI contrast imaging (12,15-17) in mice and quantitatively measured early/acute and late phase of efflux after intrathecal administration of [^64^Cu]Cu-albumin, the latter to assess the stationary phase of CSF-lymphatic efflux. Interestingly, on visual reading of PET, brain and its sulci were barely seen with radioactivity and instead, CSF space around spinal cord and cisterns maintained the radioactivity even at 24 hours. Quantitation of the radioactivity of subarachnoid space revealed higher retention in aged mice, of [^64^Cu]Cu-albumin accompanied by prominently sustaining activity at cervical and sacral lymph nodes. Technically, together with the adjusted least amount of intrathecal injection-volume, choice of latest time point enabled visualization of late stationary phase of efflux, eventually revealing the physiologic minute but definite decrease of CSF efflux from the brain and spinal cord to the adjacent lymph nodes. We propose that [^64^Cu]Cu-albumin CSF-lymphatic efflux PET and its quantitation a day after intrathecal injection be a good tool for evaluation of changes of CSF-lymphatic efflux in mice. We further expect that intrathecal [^64^Cu]Cu-albumin CSF-lymphatic efflux PET at late time point with lymph node imaging will be a good alternative over intrathecal contrast MRI CSF imaging (8-11) due to its use of smaller amount of injection volume and mass as contrast for physiologic/pathophysiologic studies in humans.

The brain waste materials, having been considered to contribute to the brain aging physiologically or pathologically (i.e. hemorrhage, neuroinflammation, trauma or neurodegenerative diseases), are first cleared from the brain parenchyma into CSF within cerebral ventricles and subarachnoid space (21,32,35-37), and then disposed through the cranial and spinal meningeal lymphatic vessels to the systemic lymph nodes (1,2,6,12,15,16,19,23,27,35,38,39). This distinctive clearance process, running along the entire length of skull and spine, is called as lymphatic efflux of CSF. As meningeal lymphatics of brain/spinal cords were linked with peri-cranial/vertebral lymph nodes, both cervical (16,19,23) and sacral (27,38), pathways of efflux of CSF were easily assumed to be separate from brain or spinal cord segments to the systemic lymphatics. Two noteworthy points are 1) wastes can take paths via whichever easier and shorter routes to reach outside of brain and spinal cords and 2) this path can also function as neuroimmune interface between central immune system (i.e. bone marrow) and central nervous system (brain and spinal cords) (3,4). It remains to be proven whether the flow from CSF to meningeal lymphatics is literally segmental, that is to say, whether vertebral segmental routes are taken separately by lymphatic channels in vertebral segmental divisions. According to the recent two reports, two possibilities are there. One is, just like brain lymphatics from nasal cavity to reach deep cervical lymph nodes (as secondary stop) via superficial cervical lymph nodes or 2) brain meningeal lymphatics directly to deep cervical lymph nodes (as primary stop), the vertebral lymphatics can come out from vertebral segmental subarachnoid space to sacral lymph node as a primary stop to reach iliac lymph node as secondary stop (27). The other is, every vertebra has its epidural lymphatics to gather to the smaller paravertebral lymph nodes, as was implied by mouse whole-body vDISCO study with transparent mounting/imaging (30). At least a long column of many vertebral segments with epidural lymphatics would have connection with the dura outlets and would make primary lymph nodes in their viscinity, which take the role of gates for adaptive immune surveillance against brain/spinal cord-derived waste and debris (38). A recent report was surprising in that meningeal lymphatics were the route for senescent astrocytes to be deployed to the systemic immune system (39).

In addition to the decreased and delayed efflux of CSF to the lymphatics in aged mice, we could also visualize clearly the lymph nodes, both cervical and sacral. This convince the physiologic meaning of the inter-communication between brain/spinal cord parenchyma and the regional lymph nodes. Then question arises about how these lymph nodes communicate with the central immune system, i.e. bone marrow of skull and vertebrae. Interestingly, Cai et al. discovered the venous connection between bone marrow sinus of the skull and venous sinus of the dura (40), which would act as a good connection path between these two central systems, neuro and immune, of brain/spinal cord parenchyma and bone marrow of skull/vertebrae. Now, we know that myeloid/lymphoid cells commute to-and-fro between the bone marrows of the skull and vertebrae and brain parenchyma via CSF (3,4). Lymph nodes are the intermediary station of the information contained in the macromolecules or cells from CSF per every level of brain and spinal cord segments. Kwon et al. (16) and Ma et al. (27) reported the sacral efflux path explicitly in animals and Jakob et al. (38) visualized the epidural lymphatic channels of vertebrae, which convey CSF solutes to regional lymph nodes (39). In addition to the nasal routes and the meningeal lymphatics to the superficial and deep cervical lymph nodes, the paravertebral epidural lymphatic channels to the nearby lymph nodes were added for every vertebral segments. In our study of [^64^Cu]Cu-albumin using PET, segmental CSF-lymphatic efflux was taken in a snapshot. Via these pathways, macromolecular solutes and senescent/injured brain/spinal cord cells can meet innate/adaptive immunity of the bodily immune system. Further visualization study for the detailed kinetics of CSF contents of this segmental communication is warranted. This will disclose the meaning of the closed loop of immune surveillance of brain and spinal cord and regional lymph nodes.

For easy visualization and quantification of the physiologic process of the body, CSF/lymphatic efflux PET using [^64^Cu]Cu-albumin provides advantages over other imaging modalities. Other radionuclide imaging studies used to show a relatively high background because we administer radionuclide-labelled radiopharmaceuticals intravenously, however, in this study, we administered the tracer intrathecally, i.e. within a compartment. Intrathecal administration can show the best contrast as was well recognized in clinical radioisotope voiding cystography having been used decades long. No radioactivity is anywhere in the body, especially in the vicinity, before the intrathecal injection of [^64^Cu]Cu-albumin. A high signal-to-noise ratio of intrathecal radioisotope-labelled albumin PET cisternography renders the versatility of qualitative and quantitative assessment of small structures such as deep cervical, sacral, or iliac lymph nodes. Interesting was the finding that hot radioactivity in superficial cervical lymph nodes reflecting the lymphatic drainage through the nasal cavity from CSF (6,7,17,41) and the sustained ratio of activities of superficial cervical lymph nodes and deep cervical lymph nodes along 9 min, 1/2/4/6/24 hrs after injection. Cervical lymphatic tracts connect superficial and deep cervical nodes (42) and cranial meningeal lymphatic vessels can drain directly to deep cervical lymph nodes (1, 2, 6, 7). As albumin (65 kD) was used as a surrogate of extracellular (CSF) tau (ca. 60 kD) or amyloid beta (100-150 kD), the sustained radioactivity even at 24 hr image in the lymph nodes implied pathologic proteins flown out from CSF to the CNS-sentinel lymph nodes would have chance to be exposed to systemic immune surveillance, i.e., antigen-processing/presenting myeloid cells and adaptive immune cells in the lymph nodes.

Various diseases were suggested to be related with CSF/lymphatic efflux dysfunction in the literature (41,43-52). Among them, in mouse models, subarachnoid hemorrhage (SAH) (36) and intracerebral hemorrhage (44) showed drainage of erythrocytes and blood solutes as expected via CSF/meningeal lymphatic efflux paths. In contrast, subdural hematoma was either drained (45) or not (44) via meningeal lymphatics. Brain tumor was also suspected to clear their components via meningeal lymphatics to cervical lymph nodes to invoke immunosurveillance and traumatic brain injury was associated with meningeal lymphatic dysfunction (47). Neurodegenerative diseases including Alzheimer’s disease (AD) were with lymphatic drainage dysfunction and interestingly, deep cervical lymph node ligation (35) aggravated AD-like pathology (49), which was reversed by focused ultrasound (50) or theta burst electrical stimulation to the brain (45). The lymphatic ablation using chemical Visudyne (25,31,43,52) aggravated pathology of AD or SAH. Meningeal lymphatic dysfunction was supposed to be associated with VEGF-C and lymphangiogenesis (31,41,46). The subtlety and delicacy of meningeal lymphatic dysfunction mandates the meticulous examination of CSF/lymphatic efflux, when the most importantly, we need to use the smallest volume amount of intrathecal administration of contrast, regardless of the contrast materials’ nature. We worked out the appropriate material, [^64^Cu]Cu-albumin, and the volume and injection speed for intrathecal administration in this study.

For the same purpose to understand the CSF/lymphatic efflux, initial pioneering studies (1,2,14) used ex vivo fluorescent microscopy imaging in order to visualize/quantify glymphatic or paravascular flux/reflux of intra-cisterna magna injected fluorescent dyes. The investigators found size dependence of efflux from CSF (14,15) as well as influx through perivascular spaces from subarachnoid space (19), which were reported to differ between young, middle age, and old mice. Of course, young mice had deeper perivascular penetration of intra-cisterna magna injected sizable fluorescent dye-labelled dextran (3 kD) or -ovalbumin (45 kD) (19). These were repeatedly reproduced by multi-institute study reported by Mestre et al. (53) but challenged by Smith et al. about the mechanistic interpretation of the observed findings (54). No matter how we agree with these competing ideas of the convection/advection by glymphatic mechanism or of the diffusion across any interstitial fluid space, perivascular space is patent in physiology between basement membranes of glia limitans and capillary endothelium. It conveys waste/debris from brain/spinal cord parenchymal interstitial fluid space to the CSF. Intrathecal gadolinium contrast, mostly gadobutrol (450 kD), studies in mice or rats disclosed even the subtle difference of influx to perivascular space to yield the effects on this glymphatic flows in anesthesia or sleep (22,23,26,55-58). However, unlike initial fluorescent microscopic studies, not so profound perivascular efflux was found in these MRI experiments, but only the shallow infiltration especially in rats (55-58).

Intra-cisterna magna injection (despite confirmed normal intracranial pressure on monitoring) versus intrathecal injection and also larger fractional amount of contrast (relatively in mouse) versus smaller fractional amount of contrast (in rats) and larger quantity of fluorescent contrast in ex vivo/in vivo fluorescent microscopy versus relatively smaller quantity of contrast (450 kD gadobutrol with smaller molecular weight) in MRI compared to fluorescent microscopy were the major confounders of these studies. We speculated the volume amount was the primary confounder and as we saw no stay in case of DTPA labelled with ^99m^Tc, albumin (65 kD) with trace amount was the optimal candidate for most physiologic examination. And as we did not see exactly the brain parenchymal perivascular influx, we gave up the evaluation of glymphatic especially in this investigation of age-related decline of glymphatic or interstitial to CSF efflux, which is measured by reversing the physiologic flux by the administration intra-cisterna magna with larger volume amount (10 to 15 μL) with tolerable but high speed (1 to 5 μL/min) in previous studies (**Supplementary Table 2**). We tried to decrease the volume as much as possible and with 0.3 μL /min speed, we observed the stagnation at the intrathecal space at the injection site, not moving on to the basal cistern and thus not riding the tide of CSF-lymphatic efflux to reach lymph nodes. As previously described, in mice 0.7 μL/min to infuse a total amount of 6 μL was our optimal (**Supplementary Figure 2**) (18).

There is still an ambiguity in understanding the flow of lymphatic from primary lymph nodes to secondary lymph nodes. As above, which explained the relation between superficial and deep cervical lymph nodes, sacral lymph nodes and iliac lymph nodes appeared at a similar time after intrathecal administration. Sacral lymph nodes drain sacral epidural lymphatics, however, contrary to the interpretation and suggestion by Ma et al. that sacral spine CSF flew out to the sacral lymph node and iliac lymph node collected lymph solely flowing out from sacral lymph nodes, we suspect that iliac lymph nodes drain the CSF/lymphatic efflux from spinal dura meningeal lymphatic vessels as well as lymph flow from sacral and the lower (or lumbar) vertebrae. Along 9 min, 1/2/4/6/24 hours of imaging points, at 1 hr images at earliest the iliac lymph node was visualized with sacral nodes. We consider that the iliac lymph node is the collecting site of the vertebral segmental lymphatic efflux from spinal dura meningeal lymphatic vessels including efferent lymph from sacral lymph nodes. This makes sense that the recent bidirectional communication between the systemic immune system and central nervous system, i.e. brain and spinal cord. Hosang et al., recently reported that infectious inflammation of the lungs affected the states of microglia of the thoracic spinal cord (59). The compartmentalized composition between bodily immune structures (lymph nodes) and spinal cord segments and brain might be the platform/frame of the understanding periphery to central nervous system interaction, especially gut with microbiome and brain/spinal cord (60,61). We propose [^64^Cu]Cu-albumin PET be used to scrutinize noninvasively as possible, the kinetic of CSF-lymphatic efflux easily till 24 hours after the intrathecal administration, without further manipulation but with repeated imaging. This method revealed that aging-related decline was there in normal aging mice.

Cause and effect relationship and its therapeutic significance should be tested by physiologic repeated imaging of small animals using sophisticated PET/CT or PET/MRI. Radiopharmaceutical use allows so small quantity to avoid any concern of toxicity or perturbation of stationary state of animals for observation. Animal models of traumatic brain injury, intracerebral/subarachnoid hemorrhage or even aging/sleep/circadian cycles are to be accompanied by diversity of the degree and severity or time points of model disease course in individual animals. Neuroinflammation or dementia models are also to be observed repeatedly during their course of disease even at 3/7/12/18 months of age. For this sort of studies at multiple time points., [^64^Cu]Cu-albumin PET study is recommended, in which experiment, if one is interested the normalcy or dysfunction of CSF/lymphatic efflux as the major candidate parameter of disease and/or effect of intervention, the clearance halve-lives as well as the surrogate of retention, the %IDs at 4, 6 or 24-hours post injection, is recommended as a group of imaging biomarkers for evaluation of lymphatic efflux of CSF. This is because the significantly longer clearance half-life and higher retention of tracer in aged mice was concordant with the existing evidence for age related phenotypic change that suggested a dysfunctional clearance of brain waste (25). Biomarkers along with the corresponding protocol can be applied for assessment of response to novel therapeutic strategies that are known to enhance the lymphatic efflux (50-52,62).

## CONCLUSIONS

A significantly higher clearance half-life along with significantly higher retention of tracer within the subarachnoid space was found in the aged animal. The imaging parameters of the PET protocol used in this study can be applied for the evaluation of lymphatic dysfunction as well as the assessment of response after therapeutic enhancement of lymphatic function.

## Supplementary Notes

**Supplementary Note 1.**
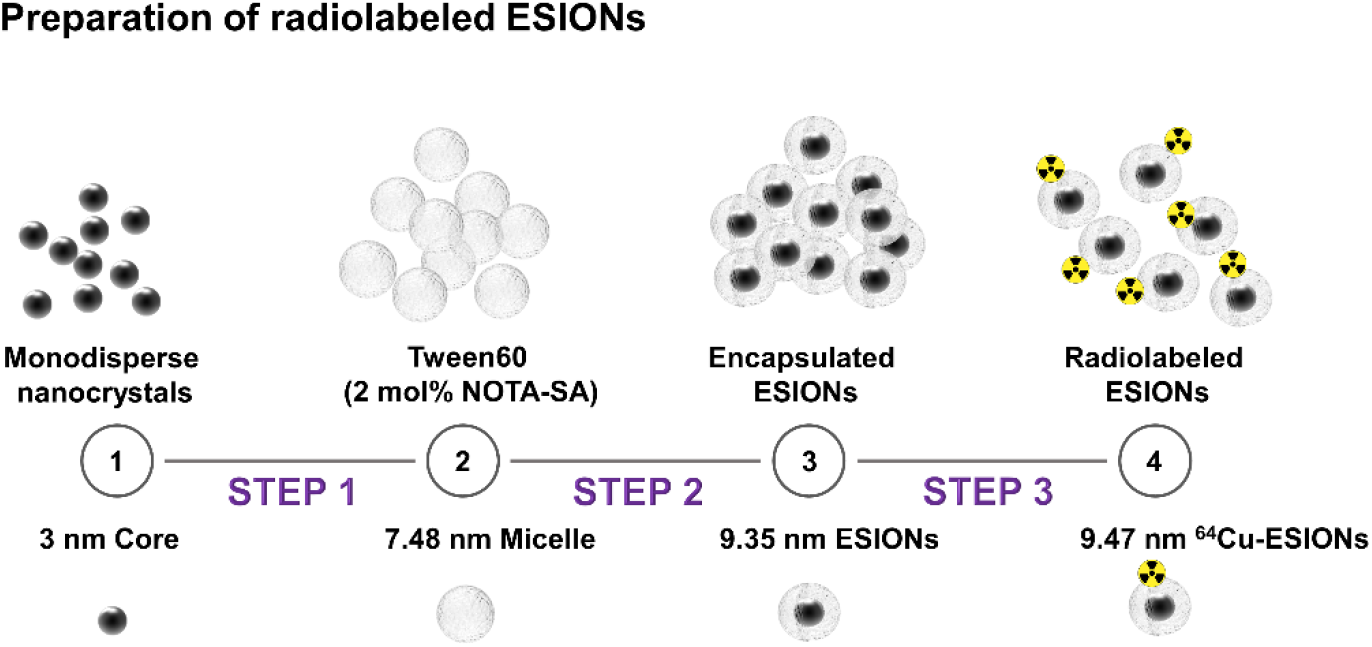
Labeling of [^99m^Tc]Tc-DTPA and [^64^Cu]Cu-ESION (extremely small iron oxide nanoparticles) ^99m^Tc was labeled with DTPA according to the routine protocol (18). ESION was labeled with ^64^Cu to make [^64^Cu]Cu-ESION according to the in-house established protocol (63). Briefly, ESIONs were monodispersed using Tween 60 with NOTA-SA and sonication. After purification of the encapsulated dispersed NOTA-surface decorated ESIONs, ^64^Cu was labelled with encapsulated EISON for their surface NOTA. Radiolabeling efficiency was 95%< and stable in vitro for one day.

**Supplementary Note 2. Imaging of intrathecal [**^**99m**^**Tc]Tc-DTPA SPECT and intrathecal [**^**64**^**Cu]Cu-ESION (extremely small iron oxide nanoparticles) PET**

Intrathecal [^99m^Tc]Tc-DTPA single photon emission computed tomography (SPECT) was done after intrathecal injection of 2 μl/min for 10 minutes (18). [^99m^Tc]Tc-DTPA SPECT was done using NanoSPECT/CT plus (Mediso). Total 24 projections into an 80 × 80 acquisition matrix were obtained with the frame time being 15 seconds for SPECT acquisition. The SPECT reconstruction used a 3-dimensional ordered-subsets expectation maximum (OSEM) algorithm. [^99m^Tc]Tc-DTPA SPECT showed CSF clearance to the systemic circulation and urinary excretion but did not show any lymph node activity along the paravertebral areas (18). Neither was also the case with superficial or deep cervical lymph nodes. We concluded that [^99m^Tc]Tc-DTPA (or similar [^64^Cu]Cu-NOTA) would not be an appropriate agent for CSF-lymphatic efflux SPECT or PET studies because they showed the clearance of CSF along time, but no evidence that clearance was vis lymphatic channels and lymph nodes.

**Supplementary Table 1.** Clearance parameters of individual mice after intrathecal [^64^Cu]Cu-albumin administration from subarachnoid space

|  | Half-Life<br>(min) | Plateau<br>(%ID) | K (min<sup>-1</sup>) |
| --- | --- | --- | --- |
| Adult mice |  |  |  |
| (n=7) | 106.6 | 15.28 | 0.006501 |
|  | 87.14 | 13.43 | 0.007954 |
|  | 72.29 | 16.16 | 0.009588 |
|  | 77.36 | 14.78 | 0.008959 |
|  | 122.2 | 14.46 | 0.00567 |
|  | 111.2 | 15.77 | 0.006232 |
|  | 77.05 | 13.78 | 0.008996 |
| Aged mice |  |  |  |
| (n=6) | 115.2 | 24.47 | 0.006017 |
|  | 141.4 | 19.55 | 0.004903 |
|  | 123.3 | 21.22 | 0.005619 |
|  | 97.97 | 20.61 | 0.007075 |
|  | 125.2 | 23.65 | 0.005536 |
|  | 136.9 | 19.63 | 0.005062 |

**Supplementary Table 2.** Infusion speed and volumes for tracer infusion in the CSF of mice reported in the literature

| Publication | Method | Route | Total volume<br>(μL) | Rate<br>(μL/min) |
| --- | --- | --- | --- | --- |
| Wang 2020 Sci Transl Med (64) | EVFM | ICM | 15 | 1.5 |
| Aspelund 2015 J Exp Med (2) | EVFM | ICM | 10 | 2 |
| Hablitz 2019 Sci Adv (55) |  |  |  |  |
| Iliff 2012 Sci Transl Med (12) | IVTPM | ICM | 10 | 2 |
| Iliff 2013 J Clin Inv (65) |  |  |  |  |
| Kress 2014 Ann Neurol (19) |  |  |  |  |
| Smith 2017 eLife (54) |  |  |  |  |
| Iliff 2014 J Neurosci (66) | EVFM | ICM | 10 | 1 |
| Xue 2020 Sci Rep (23) | MRI | ICM | 7 | 1 |
| Xie 2013 Science (67) | IVTPM | ICM | 5 | 1 |
| Achariyar 2016 Mol Neurodegener (68) |  |  |  |  |
| Louveau 2015 Nature (1) | EVFM |  |  |  |
| Ma 2017 Nat Commun (15) | NIRI |  |  |  |
| Da Mesquita 2018 Nature (5) | MRI | ICM | 2-5 | 2.5 |
| Jacob 2019 Nat Commun (38) | EVFM | ICM | 2 | 0.5 |
| Gaberel 2014 Stroke (69) | MRI | ICM | 1 | 1 |
| Wu 2018 J Nanobiotechnology (70) | IVFI | IT | 10 | Bolus |
| Jacob 2019 Nat Commun (38) | EVFM | IT | 2 and 8 | 1 |
| **This study** | **RI** | **IT** | **6** | **0.7** |

EVFM *Ex vivo* Fluorescent Microscopy, IVTP *In vivo* Two Photon Microscopy, MRI Magnetic Resonance Imaging, NIRI Near Infrared Imaging, IVFI *In vivo* Fluorescence Imaging, ICM Intra Cisterna Magna, IT Intrathecal, RI Radionuclide Imaging

**Supplementary Figure 1.**
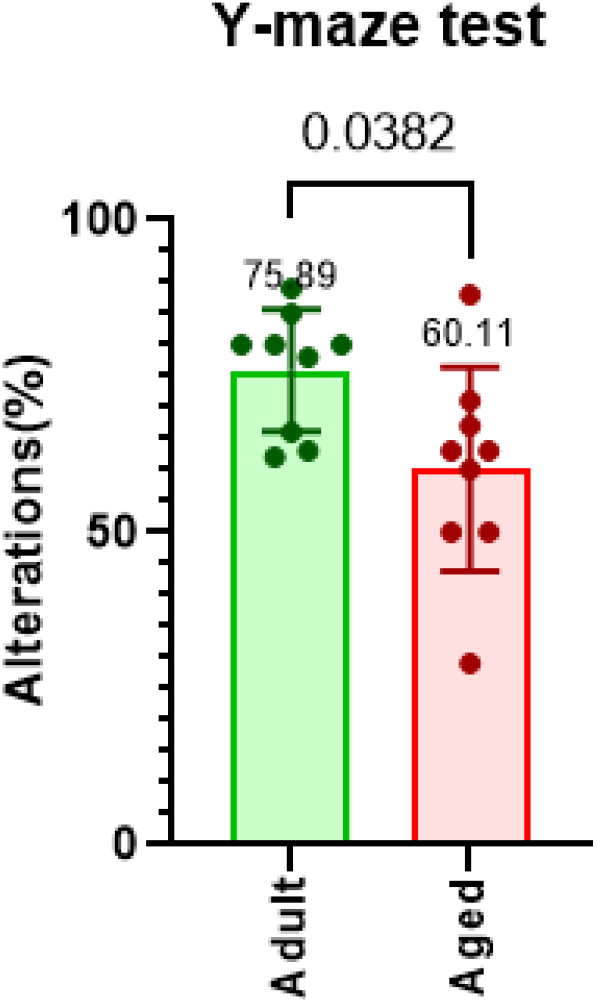
Alternation scores (%) of adult (2-9 months old) and aged mice (15-25 moths old) for spatial memory on a Y-maze test

**Supplementary Figure 2.**
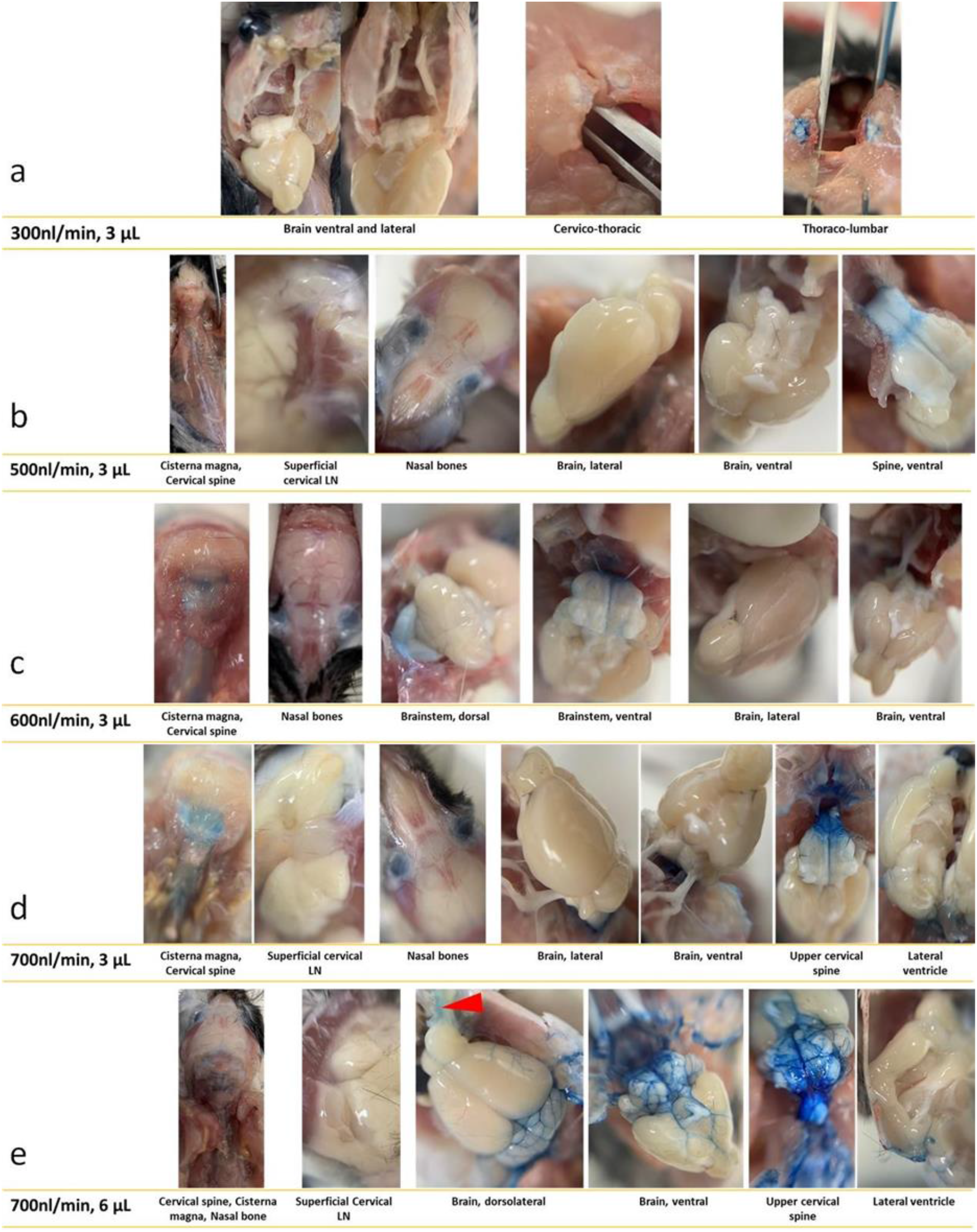
Ex-vivo distribution of Evans blue dye after intrathecal injection at different infusion speeds and volumes a) 0.3 μL/min, 3 μL, b) 0.5 μL/min, 3 μL, c) 0.6 μL/min, 3 μL, d) 0.7 μL/min, 3 μL, and e) 0.7 μL/min, 6 μL. Evans blue dye in 2% in artificial CSF was injected in adult mice; photographs were taken after cardio-perfusion with phosphate-buffered saline.

**Supplementary Figure 3.**
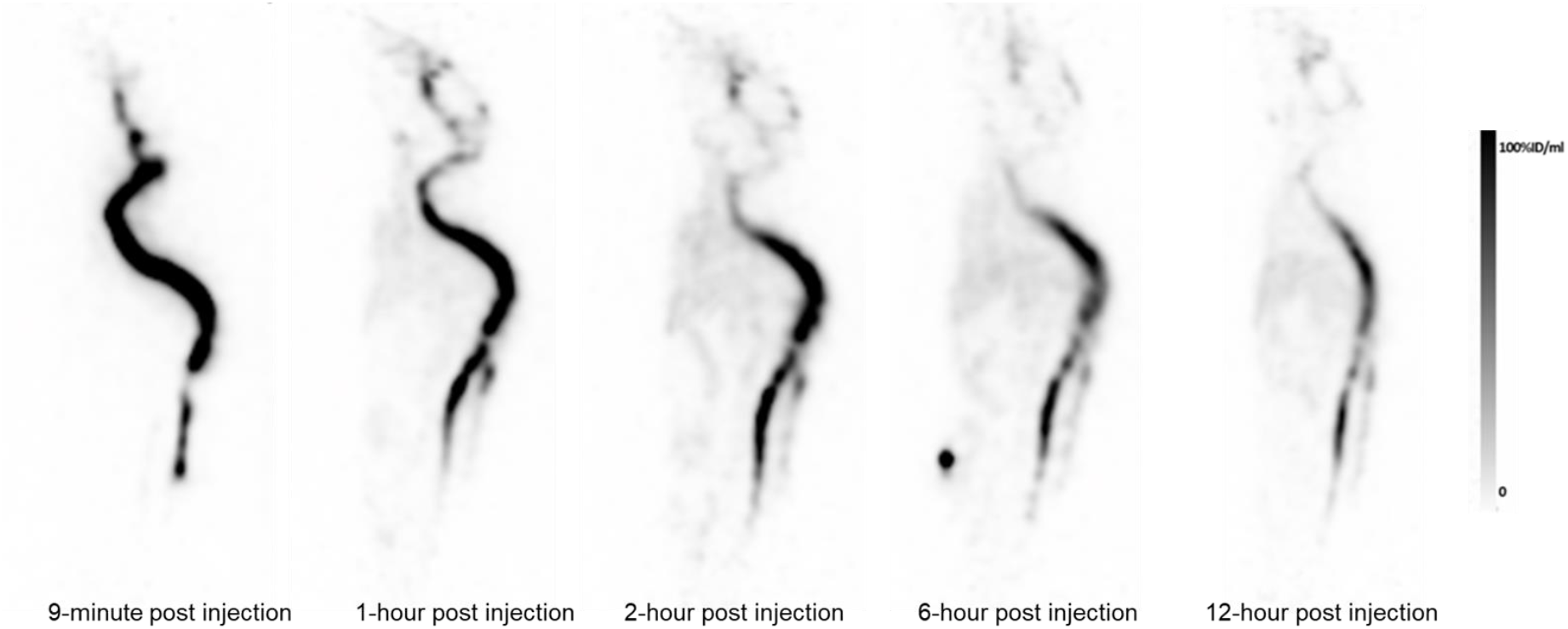
CSF/lymphatic efflux PET of intrathecal administration of microencapsulated nanoparticle [^64^Cu]Cu-ESION. This extremely small iron oxide nanoparticle, once microencapsulated and thus soluble, cleared from CSF space to the probably via lymphatics to systemic circulation. However, lymph nodes were not visualized unlike [^64^Cu]Cu-albumin. The same finding of no visualization of lymph nodes was observed with [^99m^Tc]Tc-DTPA CSF SPECT (18).

## Acknowledgments

We thank Mr. SJ Kwon and Mr. J Park for their technical assistance.

## Author contributions

AS: Methodology: Investigation: Visualization: Writing – original draft:

MS: Conceptualization: Methodology: Investigation: Visualization: Supervision: Writing – original draft: Writing – review & editing:

YC: Methodology: Investigation: Supervision:

JY Park: Methodology: Investigation:

YSL: Methodology: Investigation:

DSL: Conceptualization: Methodology: Investigation: Visualization: Funding acquisition: Project administration: Supervision: Writing – original draft: Writing – review & editing:

## Data and materials availability

All data are available in the main text or the supplementary materials. All data, code, and materials used in the analysis are available through a standard material transfer agreement with Seoul National University to academic and nonprofit by contacting the authors.

## List of Supplementary Materials

**Supplementary Movie 1**. Whole body PET maximum intensity projection images after intrathecal injection of [^64^Cu]Cu-albumin showing the organs involved in the bio-distribution of tracer.

**Supplementary Movie 2**. Whole body PET maximum intensity projection images after intrathecal injection of [^64^Cu]Cu-albumin showing the temporal change of biodistribution in an adult mouse (A), and an aged mouse (B).

